# Exploring the influence of atmospheric CO2 and O2 levels on the utility of nitrogen isotopes as proxy for biological N2 fixation

**DOI:** 10.1101/2024.03.28.587259

**Authors:** Nicola Wannicke, Eva E. Stüeken, Thorsten Bauersachs, Michelle M. Gehringer

**Affiliations:** Leibniz Institute for Plasma Science and Technology e.V., 17489 Greifswald, Germany; School of Earth & Environmental Sciences, University of St Andrews, St Andrews, Fife, KY16 9TS, United Kingdom; Institute of Organic Biochemistry in Geo-Systems, RWTH Aachen University, 52064 Aachen, Germany; Department of Microbiology, University of Kaiserslautern-Landau (RPTU), 67663 Kaiserslautern, Germany

**Keywords:** nitrogen isotopic fractionation, biological nitrogen fixation, cyanobacteria, carbon:nitrogen ratios, nodularin

## Abstract

Biological N_2_ fixation (BNF) can be traced to the Archean, over 3 Bya. The nitrogen isotopic fractionation composition (δ^15^N) of sedimentary rocks is commonly used to reconstruct the presence of diazotrophic ecosystems in the past. While δ^15^N has been calibrated under modern environmental conditions; it has not under Archean conditions, when atmospheric *p*O_2_ was lower and *p*CO_2_ was higher than today. Here we explore δ^15^N signatures in the laboratory under three simulated atmospheres with (i) elevated CO_2_ and no O_2_, (ii) present day CO_2_ and O_2_ and (iii) elevated CO_2_ and present day O_2_, in marine and freshwater, heterocytous cyanobacteria. Additionally, we augment our data set with literature data to examine for more generalized dependencies of δ^15^N during BNF across the Archaea and Bacteria, including cyanobacteria, and habitats. We find a mean ε-value of -1.38 ± 0.95, for all bacteria, including cyanobacteria, across all tested conditions. The expanded data set reveal correlations of isotopic fractionation of BNF with CO_2_ concentrations, toxin production and light, although within 1 ‰. Moreover, correlation showed significant dependency of the magnitude of ε to species type, C/N ratios and toxin production in heterocytous cyanobacteria, albeit it within a small range (-1.44 ± 0.89). We therefore conclude that δ^15^N is likely robust when applied to the Archean, stressing the strong cyanobacterial bias. Interestingly, the increased fractionation (lower ε) observed in the toxin producing *Nodularia* and *Nostoc* spp. suggests a heretofore unknown role of toxins in modulating nitrogen isotopic signals that warrants further investigation.

**Importance:** Nitrogen is an essential element of life on Earth, however, despite its abundance it is not biologically accessible. Biological nitrogen fixation is an essential process whereby microbes fix N_2_ into biologically usable NH_3_. During this process, the enzyme nitrogenase preferentially uses light ^14^N, resulting in ^15^N depleted biomass. This signature can be traced back in time in sediments on Earth, and possibly other planets. In this paper, we explore the influence of pO_2_ and pCO_2_ on this fractionation signal. We find the signal is stable, especially for the primary producers, cyanobacteria, with correlations to CO*_2_*, light and toxin producing status, within a small range. Unexpectedly, we identified higher fractionation signals in toxin producing *Nodularia* and *Nostoc* species, that offers insight into why some organisms produce these N-rich toxic secondary metabolites.

## Introduction

Biological N_2_ fixation (BNF) is a fundamental metabolism that controls the size and diversity of Earth’s biosphere. Only a subset of Bacteria and Archaea, so-called diazotrophs, possess the needed enzymatic machinery to break the triple-bond of the N_2_ molecule and convert it into bioavailable ammonia (Alberty, 2005). While several abiotic sources of bioavailable nitrogen have been identified, including lightning discharge (Barth et al., 2023; Navarro-González et al., 1998; Wong et al., 2017), volcanism (Mather et al., 2004), photolysis (Tian et al., 2011) and hydrothermal reduction reactions (Brandes et al., 1998; Rimmer & Shorttle, 2019; Schoonen & Xu, 2001), the size of these fluxes is poorly constrained and would likely have been heterogeneous over space and time. The origin of diazotrophy may therefore have been a crucial event in the early history of life that made the biosphere less dependent on abiotic planetary processes (Lyons et al., 2024; Parsons et al., 2021; Garcia et al., 2020; Mus et al., 2019, Schneider et al., 1997; Stal, 2015). On a cellular level, the enzyme nitrogenase catalyses the progressive reduction of molecular N_2_ to ammonia (NH^4+^). The most common, and generally considered most efficient nitrogenase isoform (Harwood, 2020; Luxem et al., 2020; Mus et al., 2018), the MoFe-bearing nitrogenase, uses eight electrons to convert one molecule of N_2_ into two molecules of NH_3_ and one molecule of H_2_ (Hinnemann & Nørskov, 2006; Stal, 2015):

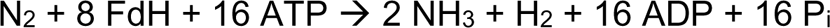

As indicated in the equation above, adenosine tri-phosphate (ATP) is involved in the process to overcome the large activation energy of BNF, being converted to adenosine di-phosphate (ADP) and orthophosphate (P_i_). ATP is then regenerated elsewhere in the cell, which, in the case of oxygenic phototrophs, occurs largely during daytime photosynthesis. All nitrogenases generate H_2_ during BNF, with the alternative isoforms of the iron only (FeFe-) and vanadium-iron (VFe-) bearing nitrogenases generating more H_2_ relative to NH_4_^+^, making them less efficient for N_2_ fixation, while placing additional energy requirements on the cell (Harwood, 2020, Hinnemann & Nørskov, 2006; Luxem et al., 2020; Schneider et al., 1997; Stal, 2015).

Earlier phylogenetic reconstructions suggested that the nitrogenase enzyme, which catalyzes N_2_ reduction to NH_4_^+^, emerged relatively late, after the Great Oxidation Event (GOE) in the Paleoproterozoic (Boyd et al., 2011; Mcglynn et al., 2013). This finding was reconciled at the time with the observation that abundances of molybdenum (Mo) in marine shales also increased after the GOE due to enhanced oxidative weathering on land and the subsequent growth of the marine Mo reservoir (Scott et al., 2008). Hence the most common variety of nitrogenase, which requires molybdenum (Bellenger et al., 2020; Mcglynn et al., 2013), may have benefited from this geochemical transition. However, more recent phylogenetic studies with large datasets have pushed the origin of nitrogenase back to the Archean eon (Pi et al., 2022, Garcia et al., 2022; Garcia et al., 2020; Parsons et al., 2021) and possibly back to the last universal common ancestor (Weiss et al., 2016). The latter may be consistent with the idea that N_2_ reduction to NH_4_^+^ could have evolved from an abiotic process occurring on sulfide minerals to an enzymatic process driven by iron-sulfur clusters (Preiner et al., 2018). Even though Archean Mo levels were minimal, likely in the lower nM range (Johnson et al., 2021) compared to the 108 nM average in the modern ocean (Pizarro et al., 2014), they appear to have been high enough to enable the emergence of the nitrogenase enzyme (Glass et al., 2012).

To further elucidate the question of when BNF emerged, several studies have looked at nitrogen isotope ratios (δ^15^N) in ancient sedimentary rocks that preserve organic matter. The two stable isotopes of nitrogen (^15^N ≙ 0.3663% of atmospheric N atoms, ^14^N ≙ 99.6337% of atmospheric N atoms) are fractionated during many biological metabolisms that take up a form of nitrogen from the environment. Chemical reactions invariably favor the lighter over the heavier isotope (Dawson & Brooks, 2001), such that a product will be depleted in ^15^N relative to a substrate.

Culturing experiments with modern cyanobacteria – the major N_2_-fixing organisms in the modern ocean – revealed fractionations (ε = δ^15^N_biomass_ – δ^15^N_N2_, where δ^15^N = [(^15^N/^14^N)_sample_/(^15^N/^14^N)_air_ – 1] x 1000) of 0‰ to -2‰ relative to atmospheric N_2_ (Bauersachs et al., 2009; Denk et al., 2017; McRose et al., 2019; Rowell et al., 1998). Slightly larger fractionations down to -8‰ were observed in deletion mutants encoding only one of the so-called alternative nitrogenases, where either Fe or V replaced Mo in the reaction site (McRose et al., 2019; Zhang et al., 2014). In contrast, redox reactions involving NH_4_^+^ oxidation to nitrate (nitrification), nitrate reduction to N_2_ (denitrification) or dissimilatory nitrate reduction to NH^4+^ (DNRA) as well as partial assimilation of dissolved NH_4_^+^ into cells are associated with much larger fractionations, including pathways that generate mostly positive δ^15^N values (Stüeken et al., 2016). This framework underpins the use of isotope data from organic-rich rocks as a proxy for the type of nitrogen metabolism that occurred during the time of sediment deposition.

In the mid-Archean, until about 2.8 billion years ago (Ga), many data points fall within the range of -2 to +2 ‰ (Homann et al., 2018; Ossa Ossa et al., 2019; Pellerin et al., 2023; Stüeken et al., 2016), which is consistent with BNF driven by a MoFe-containing nitrogenase (Ader et al., 2016; Stüeken et al., 2015). The geochemical data thus appear to confirm the conjecture that BNF originated early in Earth’s history. The necessary Mo may have been sourced from hydrothermal fluids (Evans et al., 2023; Huston et al., 2001), from anoxic dissolution of volcanic glasses (Greaney et al., 2018), or from low levels of oxidative weathering (Planavsky et al., 2014). However, a weakness in this line of argument has always been that the geochemical data are calibrated by a small set of culturing studies, summarized in Denk et al. (2017), which focused on a few select organisms. Furthermore, these experiments were conducted under standard modern atmospheric conditions, although the Archean atmosphere was enriched in CO_2_ and depleted in O_2_ relative to today (Fig. 1; Goldblatt, 2018). To our knowledge, the effect of these variables on nitrogen isotopic fractionation during diazotrophy has so far not been tested. High CO_2_ levels may spur biological productivity (Hutchins et al., 2015; Schippers et al., 2004; Wannicke et al., 2012; Wannicke et al., 2021) and thus, perhaps increase N_2_ fixation rates, while lower O_2_ levels would make the iron-sulfur cluster in nitrogenase less vulnerable to oxidation. It is therefore conceivable that the nitrogen isotopic fractionation responds to *p*CO_2_ and/or *p*O_2_, impacting the utility of δ^15^N as a biogeochemical proxy.

**Figure 1:**
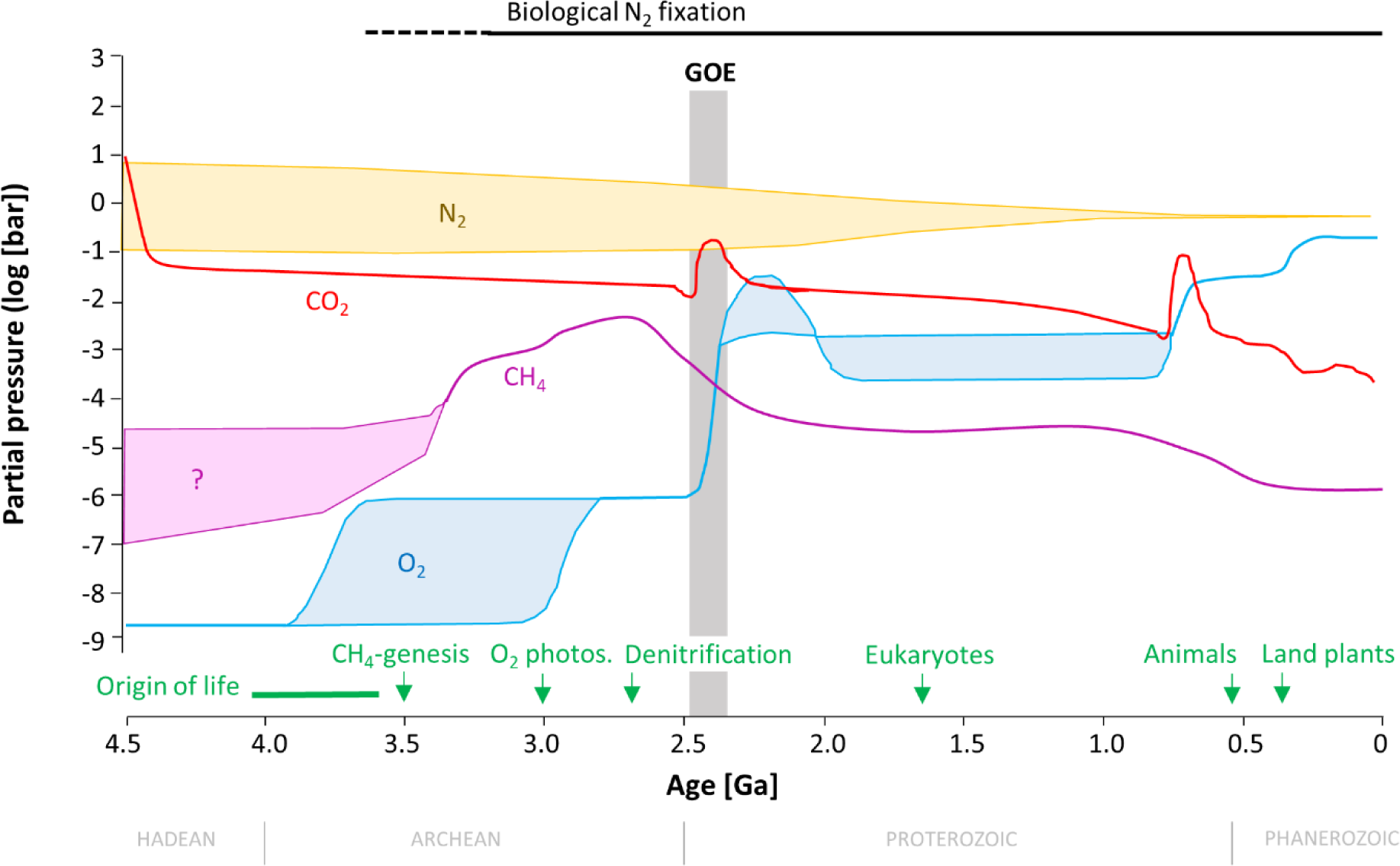
Atmospheric evolution through Earth’s history. Modified from Goldblatt (2018). Information about evolutionary events are reviewed in Lyons et al. (2024, in press). In the Archean (4-2.5 Ga), when biological N_2_ fixation evolved, *p*CO_2_ was probably two orders of magnitude higher than it is today while *p*O_2_ was at least six orders of magnitude lower.

Here we addressed some of these knowledge gaps with nitrogen isotope measurements of cyanobacterial cultures grown in the laboratory under a set of atmospheric conditions, which mimic the Archean Earth, present day and future scenarios of *p*CO_2_ and *p*O_2_. Given that cyanobacteria evolved under conditions of elevated CO_2_ when compared to present day levels, and without access to large amounts of biologically available N, we hypothesize that these organisms may respond positively to elevated CO_2_ levels and increase their N_2_ fixation rate with possible impacts of the isotopic inventory. Cyanobacteria can tightly regulate their C/N balance (i.e., the ratio of total organic carbon (TOC) to total nitrogen (TN) in the cell) (Forchhammer & Selim, 2019; Forchhammer et al., 2022), and therefore biomass C/N ratios were monitored to test if spurred CO_2_ fixation alters the stoichiometry. We further supplemented the culture data from this study with an extensive compilation of nitrogen isotopic fractionation data from other diazotrophs reported in the literature.

## Materials and methods

### a) Laboratory study

#### Strains and culture conditions

The cyanobacterial diazotrophs investigated in this study are listed in Table 1. *Calothrix* PCC7507 and *Nostoc* sp. PCC7524 were purchased from the Pasteur Culture Collection. *Nodularia spumigena* CCY 9414 was sourced from Lukas Stal (Culture Collection Yerseke, The Netherlands), *N. spumigena* NSBL06 was a gift from Hanna Mazur-Marsek (University of Gdańsk, Poland) and *Nodularia harveyana* SAG 44.85 was obtained from the Culture Collection of Algae at Göttingen University (SAG). *Nostoc* sp. 73.1 is a terrestrial cyanobacterium isolated from a coralloid root of a *Macrozamia* cycad (Gehringer et al., 2012; Gehringer et al., 2010) and has been deposited in the German Collection of Microorganisms (DSMZ) as strain (DSM114167). Brackish dwelling *Nodularia* species were cultivated in Baltic Sea medium (Ba_0_) lacking combined nitrogen sources (Wannicke et al., 2021). Terrestrial *Nostoc* and *Calothrix* species were also grown under diazotrophic conditions in standard BG11_0_ (Gehringer et al., 2010). Triplicate T_175_ ventilated cell culture flasks (Sarstedt, Germany) containing 100 ml of the appropriate medium, was inoculated with 50 ml of a fresh cyanobacterial inoculum in exponential phase. Flasks were laid flat to maximize gas exchange with the respective atmospheres and minimize shading effects. Experimental cultures were maintained under a high CO_2_ (eCO_2_) atmosphere comprising normal air supplemented to 2000 ppm CO_2_ (Plant growth chamber E-22L, Percival, USA) or an anoxic atmosphere comprising N_2_ gas supplemented with CO_2_ to 2000 ppm, denoted as AnoxHC (GS MEGA 4 Glovebox, Germany). Control cyanobacteria were grown at present day levels of CO_2_ (∼ 400 ppm) defined here as the low CO_2_ (PAL) atmosphere (Plant growth chamber E-22L, Percival, USA). All cultures were acclimated to the alternative atmospheres for at least 4 months prior to setting up cultures for stable isotope analysis. Individual strains were cultured under identical conditions at 24 ^°^C, 60% humidity and 60 µmol photons m^-2^ s^-1^ with a 10:14 h light:dark cycle. Biomass was harvested after 3-4 weeks, when the cultures had reached stationary phase, by centrifugation in sterile 50 ml Falcon tubes, washed twice with sterile MQ water and frozen at -80°C. After lyophilization for 48 h, aliquots of the dried biomass were sent for isotopic analysis.

**Table 1.** Summary of the samples generated for isotopic analysis in this study. In total, 82 samples from cultures of the diazotrophs listed below were investigated to assess the potential effect of changes in *p*O_2_ and *p*CO_2_, and culture format on δ^15^N fractionation. n.d.-not determined

| Diazotrophs |  | Culture |  | Atmosphere |  |  |
| --- | --- | --- | --- | --- | --- | --- |
| Species | Strain | Medium | Format | PAL | eCO<sub>2</sub> | AnoxHC |
| *Nostoc* sp. | 73.1<br>(DSM114167) | BG11<sub>0</sub> | liquid | n=9 | n=17 | n=4 |
|  |  | BG11<sub>0</sub> | Mat (basalt)<br>Mat (sand) | n=3<br>n=3 | n=2<br>n=3 | n.d.<br>n.d. |
| *Calothrix* sp. | PCC7507 | BG11<sub>0</sub> | liquid | n=3 | n=3 | n=5 |
| *Nostoc* sp. | PCC7524 | BG11<sub>0</sub> | liquid | n=3 | n=3 | n=5 |
| *Nodularia spumigena* | CCY9414 | BA<sub>0</sub> | liquid | n=3 | n=3 | n=3 |
| *Nodularia spumigena* | NSBL06 | BA<sub>0</sub> | liquid | n.d. | n.d. | n=4 |
| *Nodularia harveyana* | SAG 44.85 | BA<sub>0</sub> | liquid | n=3 | n=3 | n.d. |

Mats of *Nostoc* sp. 73.1 (DSM114167) were generated to test whether terrestrial cyanobacteria, with direct contact to the atmosphere, presented a different isotopic signature, compared to biomass generated in liquid medium. Beach sand (0.6-1 mm) was washed thoroughly with deionized water and sterilized overnight at 230 ^°^C. Basalt obtained from Eppelsberg Quarry was similarly treated. An 80 ml volume of sand, or basalt grains, was placed in a sterile deep Petri-dish with ventilation knobs on the lid (Sarstedt, Germany). Twenty ml of BG11_0_ medium was pipetted onto the sand/basalt to moisten it. Thereafter, 5 ml of exponentially growing culture material was gently pipetted onto the sand/basalt and the closed Petri-dishes cultured as for the liquid cultures. The mats were harvested after a month, the sand/basalt grains brushed off and the biomass dried at 60 ^°^C for 48 hours before being sent for isotopic analysis.

#### Isotopic analysis

Dry biomass was analyzed for nitrogen and carbon isotopes by flash combustion with an elemental analyser (EA) IsoLink coupled via a ConFlo IV to a MAT253 isotope ratio mass spectrometer (Thermo Fisher Scientific). The EA was operated in dual-reactor mode, where one reactor, held at 1020°C, was packed with chromium oxide granules as a combustion aid, followed by silvered cobaltous cobaltic oxide granules as a sulphur trap. The second reactor, held at 650 °C, was packed with copper wire to trap excess O_2_ and to convert NO_x_ species to N_2_ gas. It was followed by a water trap packed with magnesium perchlorate at room temperature. Ca 2-3 mg of freeze-dried, homogenized biomass was weighed into tin capsules and loaded into the autosampler of the EA, interspaced with blanks and standards. USGS-40 and USGS-41 were used for calibration; USGS-62 was used for quality control. Data are expressed in delta notation (δ [‰] = R_sample_/R_standard_ – 1). For nitrogen, R = ^15^N/^14^N, and the standard is atmospheric N_2_ gas. For carbon, R = ^13^C/^12^C, and the standard is VPDB. Average reproducibility of USGS-62 was ± 0.2‰ for δ^15^N and ± 0.1‰ for δ^13^C. To calculate the isotopic fractionation (magnitude of BNF relative to the nitrogen source), epsilon, that was used during the culturing experiments (ε = δ^15^N_biomass_ – δ^15^N_N2_), we also measured the isotopic composition of the N_2_ gas tank, using the dual inlet system of the same mass spectrometer. The gas was measured three times with eight comparisons to a reference gas in each measurement, and it was found to have a composition of -10.43 ± 0.04‰. This value was used in the calculation of ε-values following the equation shown above.

### b) Expanded data set (including literature review)

To address our research question, namely whether atmospheric CO_2_ and O_2_ levels influenced nitrogen isotopic fractionation during BNF by heterocytous cyanobacteria in liquid or mat cultures, available literature was screened, initiated by the seed publication of Denk et al. (2017) and references therein. Pre-selection was done to exclude all scientific work reporting on inorganic/organic nitrogen-based growth of cyanobacteria, as well as stable isotope tracer experiments using enriched ^15^N-N_2_. To ensure inclusion of naturally occurring organisms that came into direct contact with the respective atmospheric conditions, studies reporting on nitrogen fractionation during BNF of symbionts and/or artificially created mutants, were also excluded.

To further extend the data set with organism other than cyanobacteria, publications reporting δ^15^N of organisms capable of BNF belonging to Bacteria, other than cyanobacteria, and Archaea were included in the survey. Data extraction from incorporated publications was done manually and/or upon provision by the authors themselves, and included species and strain information, atmospheric CO_2_ and O_2_ conditions, δ^15^N values, as well as abiotic meta-data (temperature, light, salinity), and information pertaining to production of the hepatotoxin nodularin (if available). In case of deviating units, conversion to a common representation was performed (e.g. AnoxHC ≙ 0% O_2_). Calculation of ε from δ^15^N values was done as described above. The reference list of incorporated literature and tabular data are available in Supplementary Tables 2 & 3.

We did not perform any statistical analysis of the effect of different nitrogenases on BNF ^15^N fractionation, because information on the type of nitrogenase occurring in a given diazotroph is frequently not reported in the specific scientific literature. In addition, we eliminated from our compiled dataset any data from organisms where MoFe was absent, because large fractionation values have been reported for artificially created laboratory mutants of *Azotobacteria vinelandii* and *Rhodopseudomonas palustris* where the Mo based nitrogenase was deleted or inactivated (McRose et al., 2019; Zhang et al., 2014). Our study thus focuses only on organisms possessing the most commonly occurring MoFe nitrogenase, which enables us to focus on the effects of environmental parameters (rather than nitrogenase choice) on the isotopic fractionation.

### c) Statistical analysis

Spearman correlations were used to identify relationships between ε and environmental factors over different groups of organisms capable of BFN (Archaea, Bacteria, Cyanobacteria). Categorical, non-numerical parameters (toxin production, organism group) were translated to numerical groups to enable further analysis (e.g., 1= nodularin production, 0= non-nodularin production, Supplementary Tables 1, 2). Correlations were considered significant when p was ≤ 0.05. Differences in ε for toxic and non-toxic cyanobacterial strains in the laboratory, as well as extended data set were analyzed using nonparametric Wilcoxon-Mann-Whitney-Test (represented by test score T) or parametric Student’s t-test (denoted by test score t). Differences in between organism groups (Archaea, Bacteria, Cyanobacteria) and atmospheric conditions (PAL, AnoxHC, eCO_2_) were analyzed using one-way ANOVA or Kruskal-Wallis one-way ANOVA on ranks followed by Post-Hoc analysis (Dunn’s Method, Bonferroni t-test). All statistical analyses were performed in Sigma Plot 13.0 (SYSTAT, Software Inc).

## Results

### a) Laboratory study

Isotopic fractionation measured in cyanobacterial species under laboratory conditions in this study ranged between 1.24 ‰ and -2.75 ‰ (Fig. 2; Supplementary Table 1). There were significant correlations between ε values and C/N ratios (Spearman’s rho = -0.326, p= 0.00439), as well as toxin production score (Spearman’s rho = -0.514; p≤ 0.001) (Table 2, Fig. 3), while no correlation was detected between ε and C/N ratios against *p*CO_2_ and/or *p*O_2_ across the whole data set. With increasing C/N ratio, ε decreases (Fig. 3A), as well as with increasing toxin scores (Fig. 3B). Correlation of ε and nodularin production was additionally verified by comparing mean values of both groups (toxic versus non-toxic). The Mann-Whitney Rank Sum Test indicates statistically significant differences (T = 1834.000, p = <0.001, n = 32/49) with lower ε values for toxin producing strains (-1.88 ± 0.65 ‰) compared to non-toxic ones (-1.52 ± 0.35 ‰, Fig. 3B) across all *p*CO_2_ treatments.

**Figure 2.**
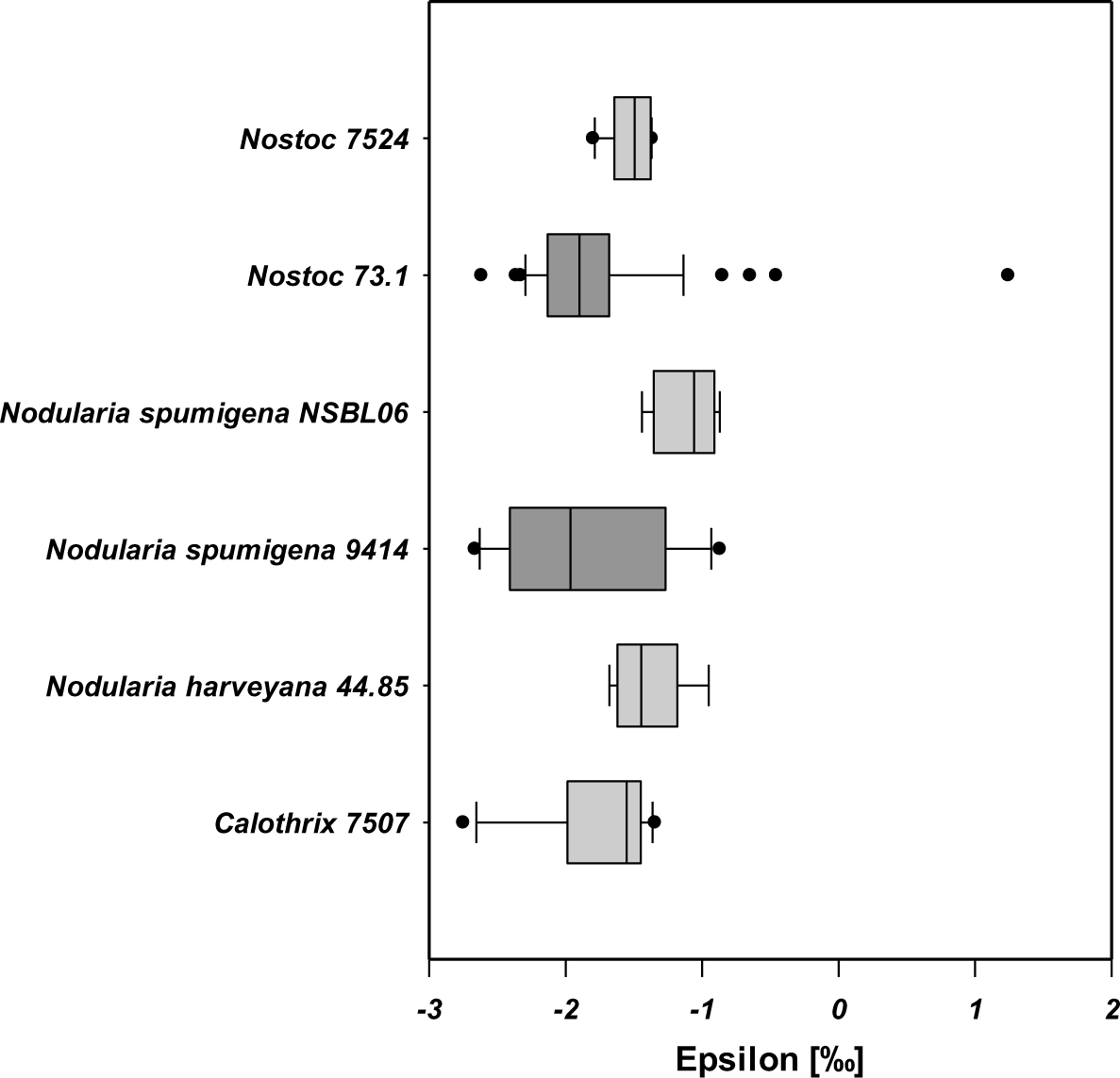
Box plots of epsilon observed in the laboratory during BNF for the different cyanobacteria investigated in this study. Dark grey box plots indicate toxin (nodularin) producing cyanobacteria, while light grey denotes non-nodularin producing strains. The spread of the box encompasses the mean 50 % of the data (=interquartile range) from the lower Q1 to the upper Q3 (25 to 75 %).

**Figure 3.**
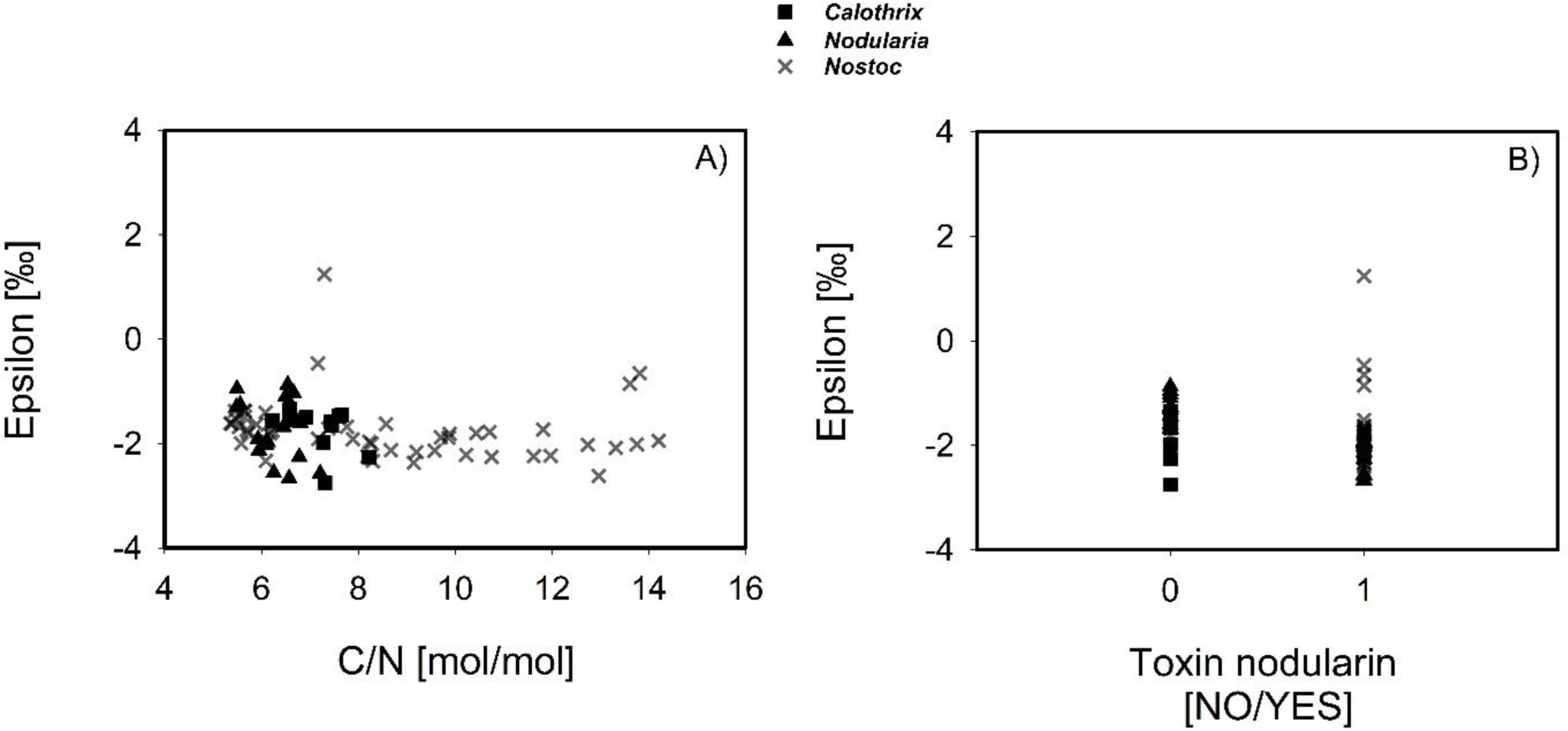
Correlation of epsilon with biomass C/N ratio and toxin profile for the species tested in the laboratory study. Values of ε displaying significant Spearman correlation coefficients from Table 2 are plotted against C/N ratio (A) and nodularin production score (B).

**Table 2:**
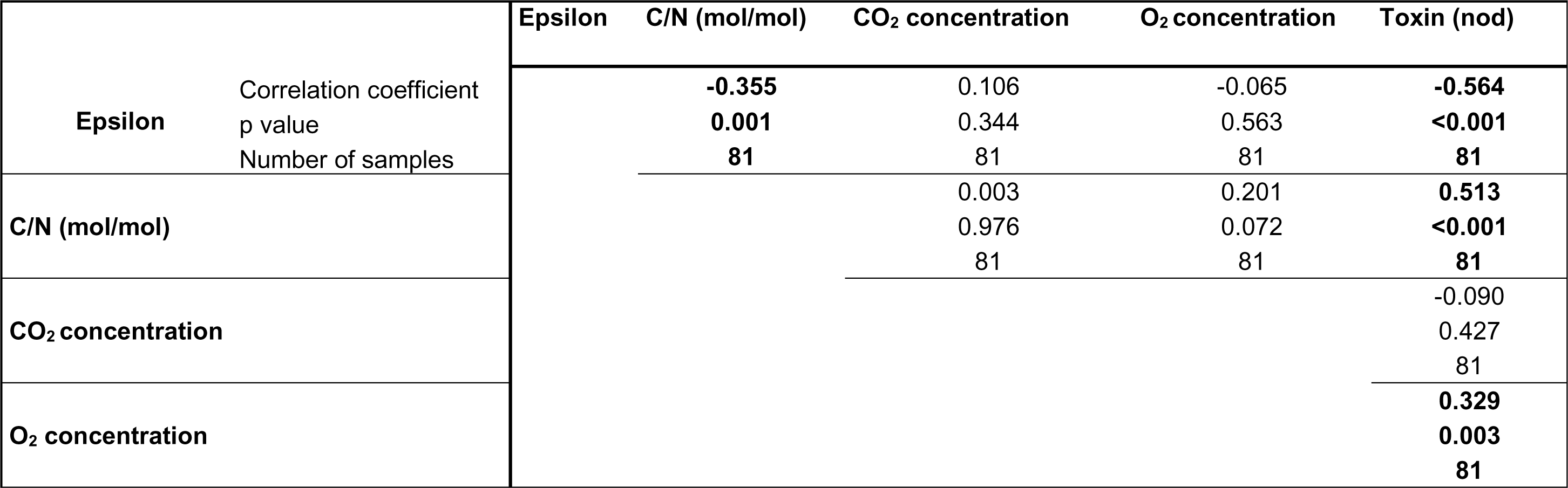
Spearman correlation of physical culture parameters vs epsilon determined in the laboratory investigation. Information on the strains is provided in Supplementary Table 1. Bold numbers indicate a significant correlation with minimum p ≤ 0.003.

Although no correlation of ε across the complete data set towards *p*CO*_2_* or *p*O*_2_* was found using Spearmańs correlation analysis, we detected differences in ε for the different *p*CO_2_ concentrations (400 ppm vs. 2000 ppm) when considering toxin and non-toxin producing cyanobacteria separately. Firstly, looking at both groups of nodularin producing cyanobacteria (toxic: *N. spumigena* CCY9414 and *Nostoc* sp. 73.1, and non-toxic: *N. spumigena* NSBL06, *N. harveyana* SAG44.85 and *Nostoc* sp. PCC7524) for which toxin status is known, significant differences in ε mean values for the two different *p*CO_2_ treatments can be found in non-toxic species (t = -3.597, p = 0.00192, n = 6/15), but not in toxin producing strains (T = 502,000, p = 0.285, n = 18/31). Secondly, when analyzing *Nodularia* and *Nostoc* species independently, opposing trends were observed. *Nodularia* displayed a dependency of ε on *p*CO_2_ in the nodularin producing strain, *N. spumigena* CCY9414 (T = 24,000, p = 0.024, n = 3/6), with lower values of ε at elevated *p*CO_2_ of 2000 ppm at AnoxHC and eCO_2_, compared to 400 ppm at PAL (-2.37 ± 0.27 ‰ vs. -1.95 ± 0.03 ‰). No significant differences in ε mean values were detected for the toxic species *Nostoc* sp. 73.1 between the *p*CO_2_ treatments (T = 309,000, p = 0.978, n = 15/25), but rather in the non-toxic *Nostoc* sp. PCC7524 (t = -3.479, p = 0.007, n = 3/8).

### b) Expanded data set

Isotopic fractionation for the two different organism-groups revealed significant differences for Archaea versus Bacteria/Cyanobacteria, with mean values of -4.13 ± 0.53 ‰ and -1.38 ± 0.95‰, respectively (p < 0.001, n = 5/256, Fig. 4). Significant correlations of ε were detected with light (Spearman’s rho = -0.176, p = 0.0197, n = 176) and *p*CO_2_ (Spearman’s ρ (rho) = -0.209, p = 0.000715, n = 261), as well as toxin production score (Spearman’s ρ = -0.436, p = 6.639E-014, n = 261; Table 3). With increasing light, *p*CO_2_ and toxin score, the ε values decrease (Fig. 5A-C). To further analyze the dependency of ε on *p*CO_2_, also in combination with toxin production, we statistically compared mean values of sub-groups of the expanded data set.

**Figure 4.**
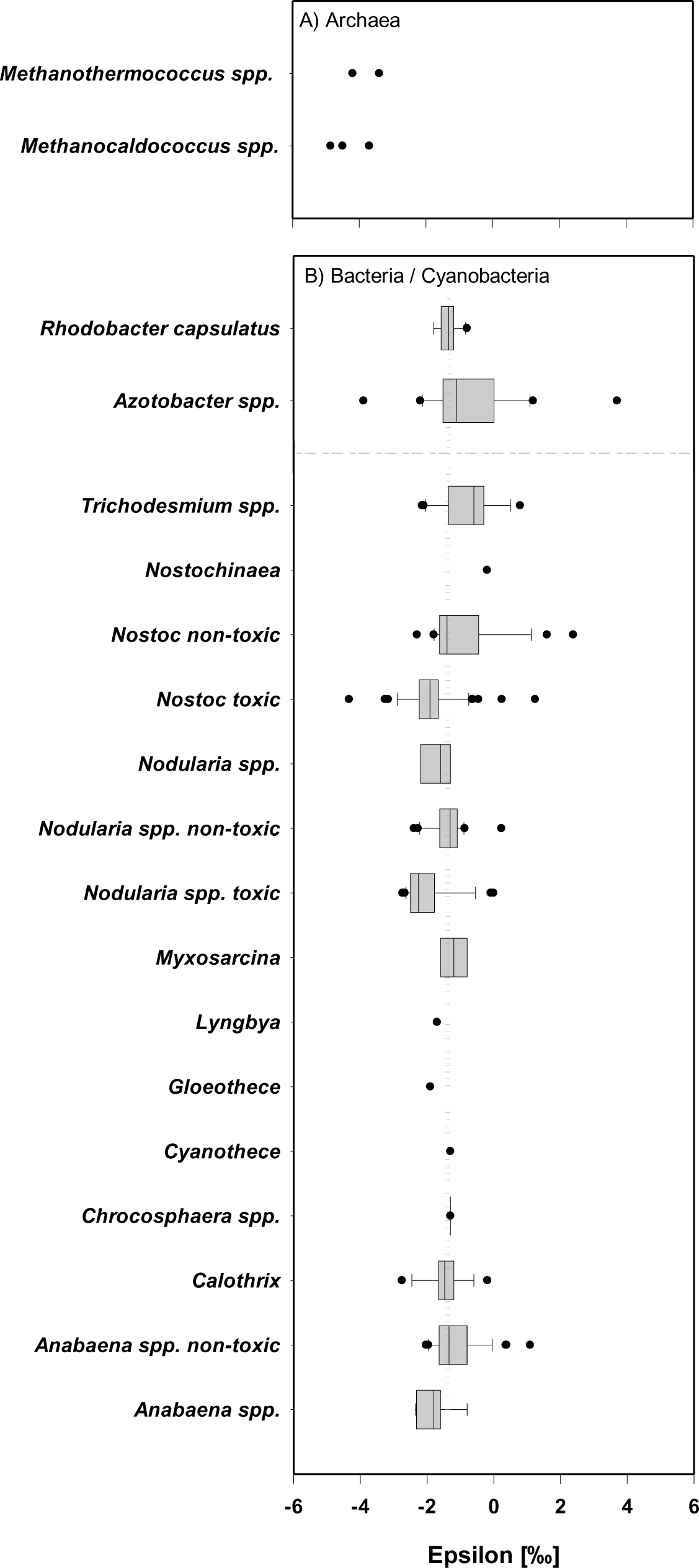
Box- and scatter plot (single data) displaying the isotopic fractionation factor (ε) for the combined set of diazotrophs. Data obtained from this study as well as literature data were categorized and plotted in three different organismal groups, namely (A) Archaea and (B) with Bacteria plotted above the dashed line, and cyanobacterial species below. Toxin-producing *Nostoc* and *Nodularia* spp. are plotted separately from known non-toxin producers. While some strains of *Anabaena* produce toxins, the toxin status of the species plotted as *Anabaena* spp. as well as *Nodularia* spp. without further specification, are unknown. *Anabaena variabilis* ATCC29413 (Thiel et al., 2014) and *Anabaena cylindrica* PCC7122 (Lyra et al., 2001) are non-toxin producers combined in *Anabaena spp.* non-toxic. For detailed information on the expanded data set see Supplementary Table 2. The grey dotted line in panel B represents the mean value of ε (-1.38 ± 0.95 ‰) across all bacterial, including cyanobacteria, data points.

**Figure 5.**
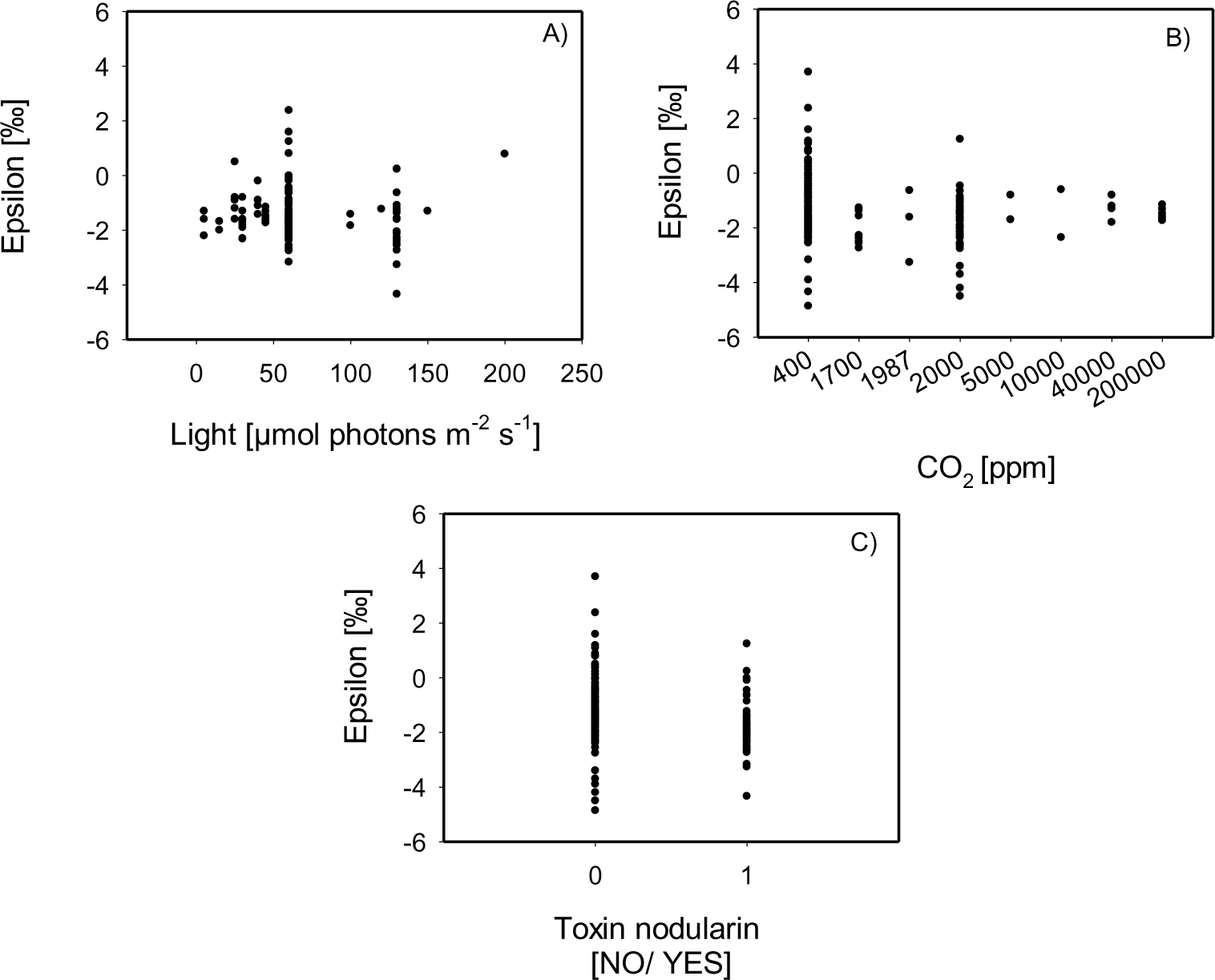
Representation of the correlation of epsilon to environmental variables in the expanded data-set. Epsilon values with significant Spearman correlation coefficients from Table 3 were plotted to highlight the correlation to light (A) and CO_2_ (B) and the toxin nodularin score (C) for the expanded dataset.

**Table 3:** Spearman correlation of parameters of the combined data sets, including literature values. Information on species and extracted data is provided in Supplementary Table 2. Bold numbers indicate a significant correlation with minimum p ≤ 0.05. C/N is not included as these values were largely not reported.

|  | Epsilon | Temperature | Salinity | Light | CO<sub>2</sub><br>concentration | O<sub>2</sub><br>concentration | Toxin (nod) |
| --- | --- | --- | --- | --- | --- | --- | --- |
| **Epsilon** | Correlation<br>coefficient<br>P value<br>Number of<br>samples | 0.122 | -0.015 | **-0.176** | **-0.203** | 0.117 | **-0.428** |
|  |  | 0.055 | 0.826 | **0.020** | **0.001** | 0.059 | **3.555E-013** |
|  |  | 247 | 215 | **176** | **261** | 261 | **261** |
| **Temperature** |  |  | **-0.296**<br>**<0.001**<br>**214** | -0.030<br>0.687<br>179 | 0.090<br>0.156<br>250 | **-0.249**<br>**<0.001**<br>**250** | **-0.281**<br>**<0.001**<br>**250** |
| **Salinity** |  |  |  | **0.290**<br>**<0.001**<br>**175** | **-01.49**<br>**0.028**<br>**218** | 0.103<br>0.128<br>218 | -0.055<br>0.432<br>218 |
| **Light** |  |  |  |  | 0.045<br>0.554<br>179 | 0.136<br>0.070<br>179 | **0.307**<br>**<0.001**<br>**179** |
| **CO<sub>2</sub>**<br>**concentration** |  |  |  |  |  | **-0.685**<br>**<0.001**<br>**264** | **0.156**<br>**0.011**<br>**264** |
| **O<sub>2</sub> concentration** |  |  |  |  |  |  | **0.141**<br>**0.022**<br>**264** |

Firstly, significantly lower ε values, within 1 ‰, were detected for the whole data set under eCO_2_ (177-2000 ppm, -1.74 ± 0.79 ‰) and AnoxHC (2000 ppm to 20%, -1.87 ± 0.91 ‰) compared to present day PAL (400 ppm, -1.21 ± 1.05 ‰) (eCO_2_ versus PAL p = 0.002, n = 48/166, AnoxHC versus PAL p = 0.013, n = 47/166, Fig. 6A, Supplementary Table 3), but overall differences in ε fall within the analytical uncertainty of 0.2 ‰. Secondly, *p*CO_2_ effects on ε of nodularin producing cyanobacteria were analyzed independently. For *Nodularia,* non-toxic strains exhibited significantly lower epsilon mean values under eCO_2_ (-2.55 ‰, single value) and PAL (-2.21 ± 0.15 ‰) conditions, when compared to AnoxHC (-1.11 ± 0.24 ‰) conditions, with t = 7.750, p < 0.001, n = 4 for PAL versus AnoxHC (Fig. 6B, Supplementary Table 3). For the toxic strain of *N*. *spumigena* CCY9414, significantly lower mean values of ε were observed under eCO_2_ (-2.48 ± 0.17 ‰) compared to PAL conditions (-1.63 ± 0.79 ‰; Fig. 6B). The mean value of ε for AnoxHC (-2.23 ± 0.23 ‰) was not significantly different from the two other *p*CO_2_ conditions (Supplementary Table 3). No significant differences in ε were recorded for *Nostoc* species under the three *p*CO_2_ conditions, regardless of toxin production score (Supplementary Table 3). Mean ε values for non-nodularin producing strains were -1.49 ± 0.11 ‰ for AnoxHC, -1.42 ± 0.05 ‰ for eCO_2_ conditions and -0.89 ± 1.23 ‰ for PAL (Fig. 6C). For toxin producing strains, mean values of ε were -1.97 ± 0.20 ‰ for AnoxHC and -1.75 ± 0.91 ‰ for eCO_2_, as well as -2.06 ± 0.88 ‰ for PAL (Fig. 6C).

**Figure 6.**
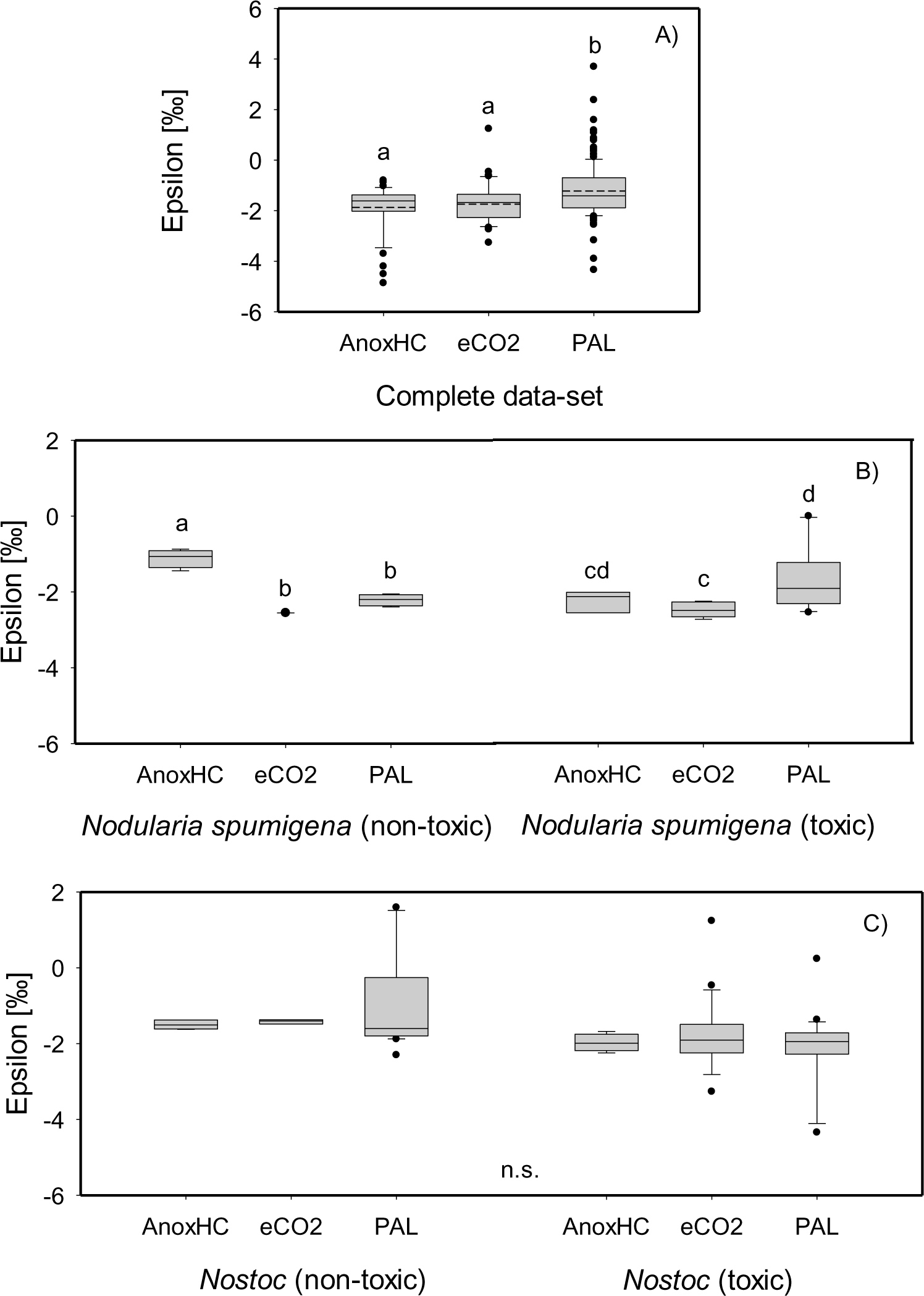
Box plot of epsilon for the different *p*CO_2_ atmospheres of the expanded data set (A) as well as for *Nodularia spumigena* (B) and *Nostoc* spp. (C) specifically. Dashed-dotted line in A) represents the mean value. Different letters denote significant differences (p < 0.05) amongst the toxin groups as determined by Duncańs post-hoc ANOVA statistical analysis, n.s.= non-significant. For detailed information on the expanded data set see Supplementary Tables 2 and 3.

## Discussion

Overall, the range of C/N and ε remains relatively narrow. In our laboratory cultures, ε values span a range of 3.99 ‰ ranging from 1.24 ‰ to -2.75 ‰. In the extended dataset, including 23 published records from the Archaea and Bacteria, along with cyanobacteria, ε spans a wide range of 8.56 ‰, ranging from 3.7 ‰ to -5.86 ‰ with a mean of -1.44 ± 0.89 ‰ for all cyanobacteria. Although we found statistically significant differences in ε in-between the atmospheric treatments, the overall differences were relatively small keeping the 0.2 ‰ analytical uncertainty of determination in mind. Some of the most negative values were recorded in MoFe-nitrogenase encoded by hyperthermophilic N_2_-fixing methanogens (Fig. 4), namely *Methanocaldococcus jannaschii* at 90 °C (Mehta & Baross, 2006), *Methanocaldococcus* at 85 °C and *Methanothermococcus* at 55 °C (Nishizawa et al., 2014). This suggests a potential albeit non-significant (p = 0.055) correlation of temperature on ε-values and motivates future investigations expanding the effect of temperature on biomass δ^15^N, beyond the 24 °C used in our study.

### What causes ^15^N/^14^N fractionation in N_2_ fixation on a cellular level?

Several studies have investigated the chemical reaction pathway of biological N_2_ reduction to NH_4_^+^ and its effects on δ^15^N, which provide an important context for our data. When Sra et al., (2004) investigated the isotopic discrimination of the Mo-nitrogenase *in vitro*, i.e., after isolation from a living organism, they measured an isotopic fractionation (ε = δ^15^N_NH3_-δ^15^N_N2_) of -17 ‰, which was interpreted as a kinetic effect. This fractionation is much larger than the values inferred from N_2_-fixing microorganisms (Supplementary Table 2, Denk et al., 2017), which led Unkovich (2013) to propose that the reduction step from N_2_ to NH_4_^+^ is not the rate-limiting step in the process. Instead, the diffusion of N_2_ into the cell and/or the utilization of the NH_4_^+^ product downstream of the nitrogenase enzyme are likely rate-limiting, such that the large kinetic effect measured by Sra et al. (2004) in an *in vitro* system is not expressed *in vivo*. This may be different in the VFe- and FeFe-based nitrogenases, the so-called alternative nitrogenases, where the reduction step is less efficient, such that it can become rate-limiting (Unkovich, 2013). This model was supported through culturing experiments with varying N_2_ partial pressures (*p*N_2_), which showed a decrease in the absolute value of ε by up to 0.6 ‰ as *p*N_2_ was changed from 0.8 bar to 0.1 bar (Silverman et al., 2019). At low *p*N_2_, the supply of N_2_ by diffusion into the heterocyte appears to become limiting, such that the isotopic fractionation associated with the reduction step has even less net impact. Limited diffusion of N_2_ and O_2_ in heterocytous strains of cyanobacteria results from thickened cell walls of the specialized nitrogen fixing cells, thereby protecting the nitrogenase from O_2,_ simultaneously also restricting diffusion of other gases such as N_2,_ CO_2_ and H_2_, a by-product of BNF (Silverman et al., 2019).

Isotopic fractionation furthermore occurs when the end-product of BNF, NH_4_^+^, is incorporated into carbon skeletons by the glutamate dehydrogenase and subsequent transfer of the amino group by a transaminase, such as glutamine synthetase/glutamate synthase (GS/GOGAT) (Flores & Herrero, 1994; Luque et al., 1994). Finally, N-containing compounds such as proteins, nucleotides, porphyrines and proteoglykanes, but also individual amino acids, may differ in respect to their δ^15^N values. The production rate of these end products ultimately depends on the growth rate of the cell and thus links the nitrogen to the carbon cycle, and ultimately to atmospheric *p*CO_2_ in the oxygenic photosynthesis in cyanobacteria. In summary, the observation that isotopic fractionations occur during N_2_-uptake into the cell and during further processing of fixed nitrogen into organic molecules motivates our search for dependencies between ε and environmental variables that impact biological N-demand.

### Is there any risk that the presence of VFe and/or FeFe nitrogenase is perturbing our data?

Examples of organisms producing either an additional VFe-based (*vnf* encoded) or FeFe-based (*anf* encoded) nitrogenase, or both, exist; however, no example has been identified to date that encodes either of these nitrogenases in the absence of the MoFe-containing nitrogenase (*nif* encoded) (Harwood, 2020, Mus et al., 2019; Bellenger et al., 2011). Additionally, the MoFe-nitrogenase is preferentially expressed under Mo replete conditions, with the VFe-nitrogenase only expressed under Mo depleted diazotrophic conditions in *A. variabilis* ATCC29413 (Thiel & Pratte, 2014) and *Azotobacter vinelandii* (Jacobson et al., 1986), the latter below 25 nM Mo, with V concentrations as low as 8 nM (Bortels, 1930; Bortels, 1940). The lowest levels of Mo are encountered in rivers, freshwater lakes and estuaries (Pizarro et al., 2014), as well as in tropical soils (Barron et al., 2009), with the modern ocean Mo levels ranging from 2-108 nM (Pizarro et al., 2014; Zerkle et al., 2006; Zerkle et al., 2008). Given the 2-3 ppm Mo in the Earth’s crust (McLennan, 2001), with some strains like *A. variabilis* able to extract Mo from silicates using metallophores (Liermann et al., 2005), BNF organisms are thought to be able to access sufficient Mo to meet their needs (Sheng et al., 2023). Mo at 10 µM and 1 µM also stimulated BNF in the archaeon, *Methanosarcina barkerei* 227, in the absence of NH_4_^+^ or glutamate (Lobo & Zinder, 1988). Collectively, these observations suggest that MoFe nitrogenase is used preferentially in biology unless organisms are under severe Mo-stress. Furthermore, ensuring that an experimental set-up is Mo-free is challenging and even cultures with just a few nM Mo have been reported to perform N_2_ fixation (Zerkle et al., 2006). Therefore, even though some of the strains investigated in this study do contain VFe and / or FeFe nitrogenase in addition to MoFe nitrogenase, we consider it unlikely that the trends in the data are impacted by the presence of these enzyme variants. This allows us to focus our discussion on the effect of environmental variables on ε and C/N ratios.

### Potential effects of *p*CO_2_ and *p*O_2_ on ε

Having established that the presence of VFe or FeFe nitrogenase does likely not impact the trends in our data, we can now turn to the investigation of environmental parameters on ε during BNF. While the effects of atmospheric N_2_ levels on ε were previously investigated (Silverman et al., 2019), the potential influence of how changes in *p*O_2_ and *p*CO_2_, as well as other environmental parameters such as light, temperature and salinity, has not.

Firstly, our study identified a significant correlation between ε and C/N content of the cultured biomass (Table 2) with decreasing ε at increasing C/N ratios. In other words, isotopic fractionations become slightly larger as the bulk biomass becomes depleted in nitrogen. This observation may indicate that incorporation of fixed N into biomass within the cell creates a minor bottleneck that imparts a small isotopic fractionation. Secondly, statistically significantly lower ε values were detected in the expanded data set at elevated *p*CO_2_ of ≥ 2000 ppm, regardless the *p*O_2_. This may be consistent with the idea that high *p*CO_2_ stimulates the growth rate in C-fixing cyanobacteria (Wannicke et al., 2021), and hence, N-demand, exacerbating the bottleneck of N incorporation into biomass. This conclusion would be consistent with the current understanding of N-isotopic fractionation during BNF described above.

Thirdly, both our laboratory investigation and the expanded literature data identified species-specific responses to atmospheric *p*O_2_ and *p*CO_2_ levels (Tables 2 and 3). Specifically, *N. spumigena* CCY9414 (marine) exhibited significantly lower ε values when grown at 2000 ppm CO_2_, irrespective of *p*O_2_, while *Nostoc* sp. PCC7524 (freshwater) showed lower isotopic fractionation (less negative ε values) under elevated CO_2_ conditions. This observation suggests that marine diazotrophs may react differently to increasing atmospheric CO_2_ levels than terrestrial or freshwater species. The difference in response between marine and freshwater organisms may be due to different environmental adaptations. Seawater is buffered by HCO_3_^-^ with a relatively constant pH leading to relatively constant CO_2_ supplies, whereas freshwater pH and dissolved CO_2_ may vary widely (e.g. McNeil & Matsumoto 2019). It is conceivable that freshwater organisms have developed strategies to maintain stability across natural environmental CO_2_ fluctuations while marine organisms lack this skill, making them more sensitive to external perturbations.

In line with this observation, the aquatic *Nodularia* species, including *N. spumigena* CCY9414, demonstrated a 6-fold higher biomass accumulation under high CO_2_ conditions when compared to terrestrial *Nostoc* species (Wannicke et al., 2021). This was accompanied by a 17-fold increase in BNF and raised PON content compared to the *Nostoc* spp., providing support for increased nitrogen demand to maintain cellular C/N ratios under high CO_2_ conditions, as recorded for *N. spumigena* sp. KAC12 (Karlberg & Wulff, 2013) and *N. spumigena* CCY9414 (Wannicke et al., 2012). C/N ratios were only slightly increased under increased CO_2_ levels in *Cyanothece* sp. ATCC51142, *N. spumigena* sp. IOW-2000/1 and *Calothrix rhizosoleniae* SC01 (Eichner et al., 2014). Species phenotypic plasticity must also be considered as *Nostoc punctiforme* CPCC41, grown at 940 ppm CO_2_ demonstrated an increase in BNF rate (Lindo & Griffith, 2017) and *Dolichospermum circinale* appeared to benefit from raised CO_2_ levels of 1700 ppm (Symes & van Ogtrop, 2019) in terms of elevated biovolume and chlorophyll a content. The recorded positive growth effects of elevated *p*CO_2_ on cyanobacteria most likely arises from down-regulation of the carbon concentrating mechanisms (Kranz et al., 2011; Levitan et al., 2010a, b and c), which are needed in most phytoplankton species to compensate for the low efficiency of the photosyntheitc enzyme ribulose-1,5-bisphosphate carboxylase oxygenase (RUBISCO) under present day CO_2_ levels (Eichner et al., 2022). The C/N ratios and ε values were, however, not determined in the above-mentioned studies.

Lastly, the lack of correlation between ε values and *p*O_2_ likely attests to the ability of diazotrophs to protect the nitrogenase enzyme from atmospheric O_2_ (Levitan et al., 2010a, b and c; Shi et al., 2010; Stal, 2015).

### Proposed impact of nodularin production on internal nitrogen fractionation

Although not within the scope of our initial hypothesis of *p*O_2_ and *p*CO_2_ affecting nitrogen isotopic fractionation during BNF, an unexpected correlation of ε to nodularin production was observed. Our laboratory investigation (Fig. 2; Table 2), identified significantly larger fractionations (more negative ε values) in the nodularin producing strains of *N. spumigena* CCY9414 and *Nostoc* sp. 73.1, compared to the non-nodularin producing strains of the same species. This observation was maintained in the expanded data set with toxin producing *Nodularia* and *Nostoc* strains having significantly lower mean ε values compared to known non-toxin producing strains of cyanobacteria (Fig. 4; Table 3). Our literature analysis suggests that some *Anabaena* species, for which toxin production is not known, may indeed be toxin producers, given their low ε values compared to the overall mean, indicating higher δ^15^N fractionation. The *Anabaena* strains of *Anabaena variabilis* (Thiel et al., 2014) and *Anabaena cylindrica* (Lyra et al., 2001) are known not to produce toxins. They also did not exhibit increased fractionation of nitrogen (Fig. 4), supporting our hypothesis that toxin production is involved in internal N-cycling, resulting in lower ε values.

The causal connection between nodularin production and increased fractionation of ^15^N is speculative so far. At least a strong correlation of the production of the N-rich (C/N ratio lower than Redfield ratio 6.6) secondary compound nodularin is undisputed, and this secondary metabolite may be removing an isotopically fractionated (^15^N-enriched) nitrogen reservoir from the bulk biomass. Nodularin is a cyclic pentapeptide with five amino acids in the peptide ring. It is the end product of a secondary metabolic pathway comprised of mixed polyketide synthases and non-ribosomal peptide synthetases (Moffitt & Neilan, 2004). The purpose of nodularin production is not fully resolved, but because of the high-energy demand of nodularin production, an ecological or physiological advantage over non-toxin producers must be assumed. A function within the cell metabolism is likely, as a rapid and covalent binding of nodularin to proteins of unknown function upon abiotic stress, was shown (Meissner et al., 2013). In addition, nodularin and microcystin (a related cyclic heptapeptide) might function as signalling or regulatory proteins (Dittmann et al., 2001). The strong connection of microcystin and nodularin turnover with the cellular N-cycle (Downing et al.,2005) is indicated by the regulation of their synthesis by the global N uptake regulator NtcA (Gehringer & Wannicke, 2014; Ginn et al., 2009), which activates transcription of nitrogen assimilation genes (Hansel et al., 2001; Tamagnini et al., 2002). The difference in cellular response to *p*CO_2_ and toxin production status is highlighted in Figs. 6B and 6C. Nodularin-producing *N. spumigena* shows a significantly larger fractionation under high CO_2_ levels, whereas the non-toxin producing strains do not. This is interesting, as aquatic *Nodularia* species overall exhibited increased BNF rates under the same *p*CO2 conditions (Wannicke et al., 2021), suggesting that toxin production has some effect on internal nitrogen cycling.

While nodularin producing *Nostoc* species also exhibited similar ε values compared to toxic *Nodularia* species, (Fig. 4; Table 3), isotopic fractionation was less affected by changing levels of *p*CO_2_. Previously, *Nostoc* spp. demonstrated significantly reduced BNF rates when compared to *Nodularia* species, under both low and high *p*CO_2_ conditions, suggesting species-specific constraints on BNF, possibly related to habitat differences (aquatic vs terrestrial) (Wannicke et al., 2021).

It is currently unknown when biological toxin production evolved. However, it is conceivable that its isotopic effect of driving the biomass to slightly more negative δ^15^N values may have impacted the geological record. This aspect is worth further investigation.

### Implications

Generally, the range in ε detected in our laboratory and expanded data-set is small compared to fractionations of well over 10 ‰ observed for biological metabolisms that involve large reservoirs of dissolved ammonium, nitrite or nitrate (Casciotti, 2009). Although there are correlations between ε, C/N ratios and *p*CO_2_ that likely relate to biological N processing within the cell, the variance in ε remains only within a few permil. Since the isotopic composition of atmospheric air has likely not changed much (< 2‰) since the early Archean (Marty et al., 2013), the ε values reported here for BNF, where the substrate is atmospheric N_2_, are equal to the δ^15^N value of biomass that would be archived in the sedimentary rock record. Comparing our range of ε values (i.e. biomass δ^15^N for diazotrophs) to sedimentary rocks over the past several billion years shows that the effects of C/N and *p*CO_2_ would likely not be resolvable (Fig. 7). Diagenetic and metamorphic alteration may additionally bias sedimentary δ^15^N values by a few permil compared to the composition of the original buried biomass, complicating the interpretation of the sedimentary δ^15^N values (Robinson et al., 2012; Thomazo & Papineau, 2013). On the contrary, the relatively low sensitivity of ε to *p*CO_2_ and C/N ratios implies that the δ^15^N signal remains a reliable indicator of BNF in deep time, at least given the current knowledge base. However, it is important to note that current data of ε during BNF remain strongly biased towards cyanobacteria. More δ^15^N information should be obtained for other nitrogenase-containing organisms including Archaea and the Bacteroides, Chlorobi, Firmicutes, Verrucomicrobia, with more representation from the Proteobacteria (Bellenger et al., 2020; Bellenger et al., 2011). As shown by the extended dataset, (hyper-)thermophilic methanogens in particular, appear to express larger ε values for BNF, suggesting that higher temperatures encountered e.g., in hot springs, may induce greater fractionation, possibly resulting from increased internal N-cycling.

**Figure 7:**
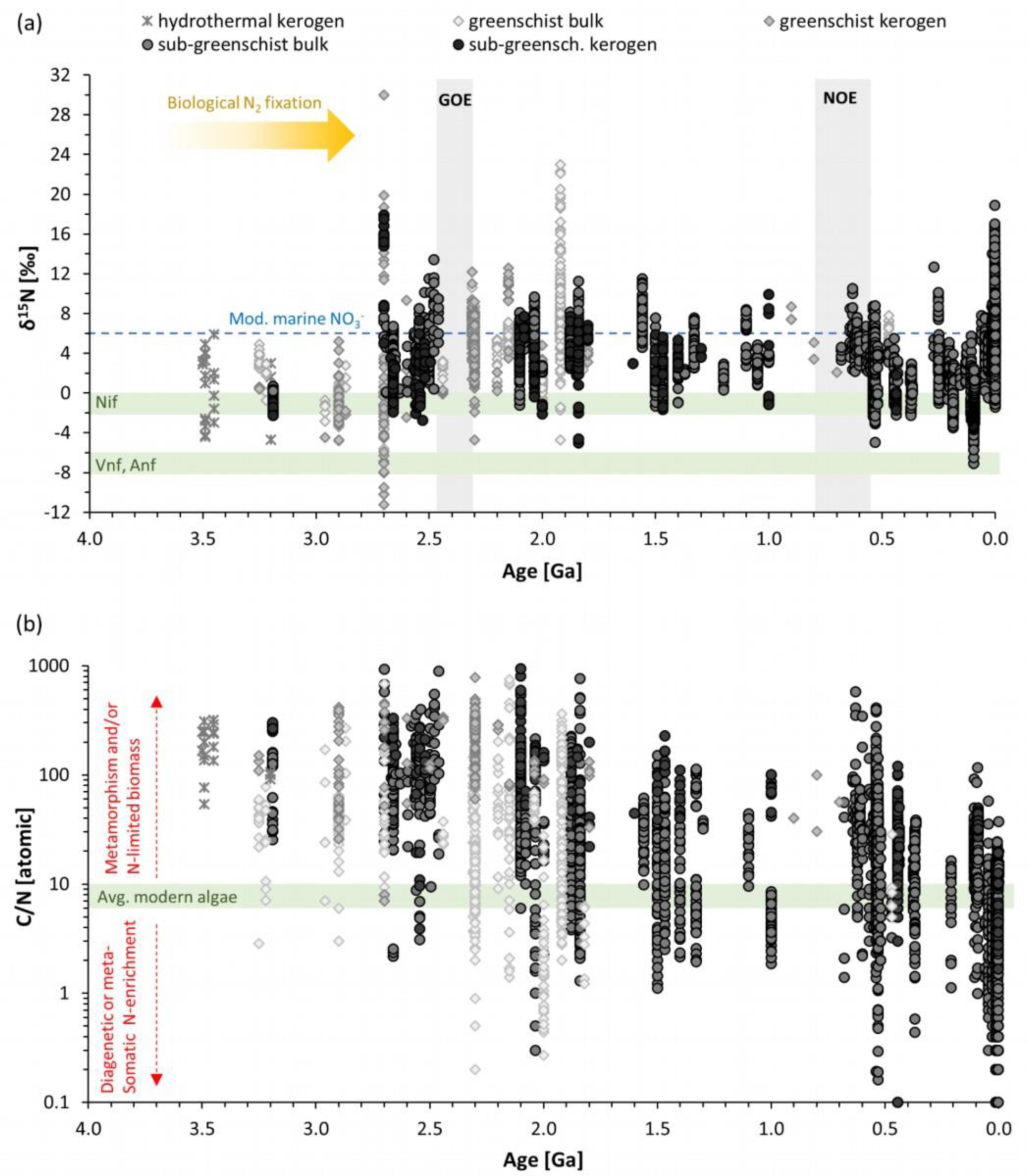
Geochemical data from the sedimentary rock record. (a) Nitrogen isotopes in bulk rocks and kerogen isolates (organic matter extracted from the bulk rock with hydrofluoric acid). (b) Ratios of organic carbon to total nitrogen. In both panels, samples are separated by metamorphic grade, and samples of amphibolite grade or higher are excluded, because such high metamorphic alteration is likely to have perturbed the primary signature by more than 2 ‰ (reviewed by Thomazo & Papineau, 2013). The data for both panels are compiled from the literature (see Johnson & Stüeken, 2024, in press).

Importantly, we emphasize the unexpected finding that toxin-producing organisms express slightly larger fractionations (more negative ε values) compared to non-toxin producers. The antiquity of toxin production is unknown, but our finding opens the intriguing possibility that the δ^15^N record may, perhaps in conjunction with phylogenetic reconstructions, provide information about the onset of this metabolic trait.

## Acknowledgements

MMG was funded by the DFG: SPP1833 grants GE2558/3-1 & GE2558/4-1. EES acknowledges funding from a NERC Frontiers grant (NE/V010824/1).

